# *Mango*: Unearthing Patterns in Large-Scale Biological Data Through Interactive Correlation Analysis

**DOI:** 10.1101/2025.10.06.680608

**Authors:** Sverre Branders, Manfred G. Grabherr, Rafi Ahmad

## Abstract

Integrating different types of biological data is often challenging due to the presence of both numerical and categorical data. This complexity makes it harder to evaluate causal biological effects, especially when confounders like population structure, sampling methods, or multi-omics integration can lead to incorrect conclusions. We introduce Mango, an interactive correlation browser designed for visually exploring any tabular data type, using a novel algorithm to correlate numerical and categorical data, regardless of their distribution, called Median-Ranked Label Encoding. Our results on genomic and transcriptomic datasets demonstrate that these correlations can effectively distinguish between biases and causal relationships in large-scale data.

## Background

As high-throughput experimental techniques become more common in life sciences, data science tools like statistics and machine learning are increasingly vital for converting large datasets into meaningful scientific insights. Relying only on automated analyses can lead to incorrect assumptions about the data. A persistent challenge in bioinformatics is ensuring datasets are free of biases. These biases—including batch effects from lab procedures or poorly stratified cohorts— are systematic and can undermine the validity of the study’s results. As a result, numerous studies have been published reporting an effect or association, but the findings do not apply to cohorts beyond those studied (1–4). Currently, there are no well-established techniques for detecting such effects in qualitative data. This emphasizes the importance of developing and validating new approaches to identify batch effects and biases, ensuring the accuracy and reproducibility of results. These effects are especially significant when, for example, examining phenotype associations in bacterial genomes, where clonal reproduction makes it difficult to distinguish causal from non-causal relationships (5). Additionally, in multi-omics – a technology that’s currently leading to the discovery of life-saving therapeutics and diagnostics – batch effects can be especially complex. This is because multiple data types with different distributions are integrated from various platforms (1). In this context, data visualization (DataVis) is increasingly recognized as a vital part of the data validation process (6). Several software initiatives exist to lower the barrier of entry and promote reproducibility in this field, including BioPerl (7), Bioconductor (8), Bioconda (9), etc. These software collections still require significant domain expertise to choose and use appropriate packages for particular biological problems. Graphical applications can further lower these barriers, making the software more accessible to researchers with different scientific backgrounds. Many domain-specific graphical software tools exist, such as EpiViz (10), which implements a Shiny-based (11) web interface that is compatible with popular Bioconductor packages.

Comparing two variables, whether discrete (such as text labels), two numeric variables, or a numeric and a discrete variable, allows for a quick assessment to determine whether the correlation is desirable or not. For example, if a clinical measurement from a sample is highly correlated with the diagnosis, then this correlation is considered desirable. Conversely, if these measurements are also highly correlated with, for example, cohorts, then this suggests that the measurements are cohort-specific rather than indicative of the disease. In the case of the latter, both cohorts and measurements will exhibit high correlation. An intuitive way to assess and explore data is through visualization.

Mango is a user-friendly standalone tool designed for exploring correlations in tabular data with minimal technical knowledge. It utilizes metadata where each row represents a sample, each column a metadata variable, and each cell an observed value. Mango addresses this by: (a) creating a correlation matrix that illustrates the similarity and redundancy among variables; (b) visually presenting these results through an interactive Graphical User Interface (GUI). This GUI enables users to access data at various levels: a high-level view featuring an interactive correlation matrix with customizable row orderings, and a detailed, zoomable view for examining individual variables.

## Results

For illustration, we used Mango on a publicly available dataset (Avershina et al. dataset) for predicting resistance to β-lactams in *Escherichia coli* based on genotype-to-phenotype analysis (12). In addition to the metadata, we analyzed the genomes of the 86 *E. coli* samples using Saguaro (13) to identify local genomic patterns, both concordant and deviant from the overall phylogeny. Saguaro produced 8 “cacti”, i.e., vectors that describe these patterns across the samples. The Saguaro cactus 0, which models 94.6% of all variable sites and is representative of the background phylogeny, is not correlated with any other variables, including the metadata and other cacti, indicating no discernible batch effects. While cacti 1 through 7 are correlated with each other, cactus 6, modeling 0.1% of all variable sites, has the highest correlations with phenotypic susceptibility (Figure 1).

**Figure 1:**
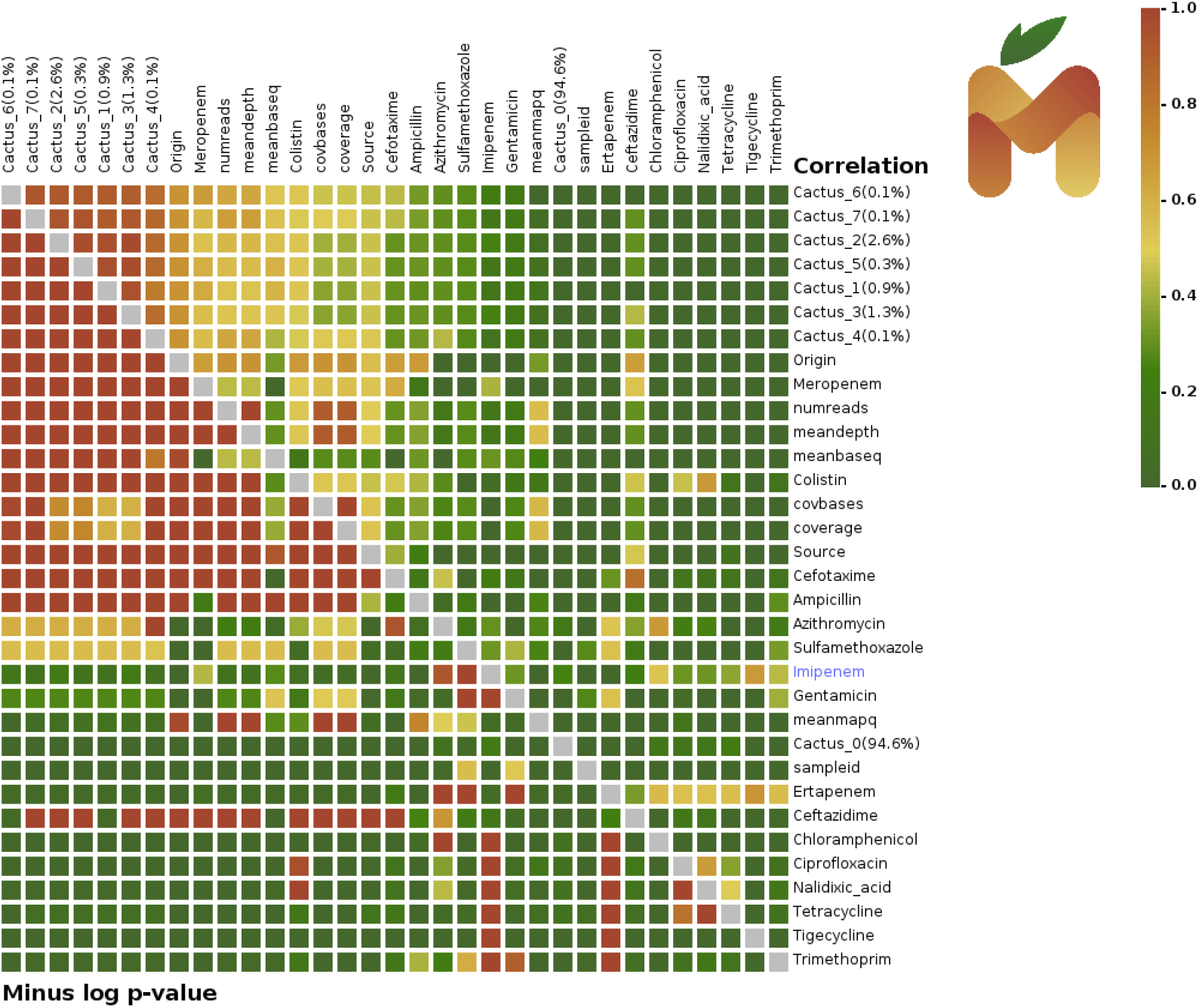
Correlation view in Mango. The matrix is sorted based on correlations to cactus 6 (top row). The top right triangle displays the correlations with color coding, while the bottom left triangle shows the p-values.

**Figure 1a:**
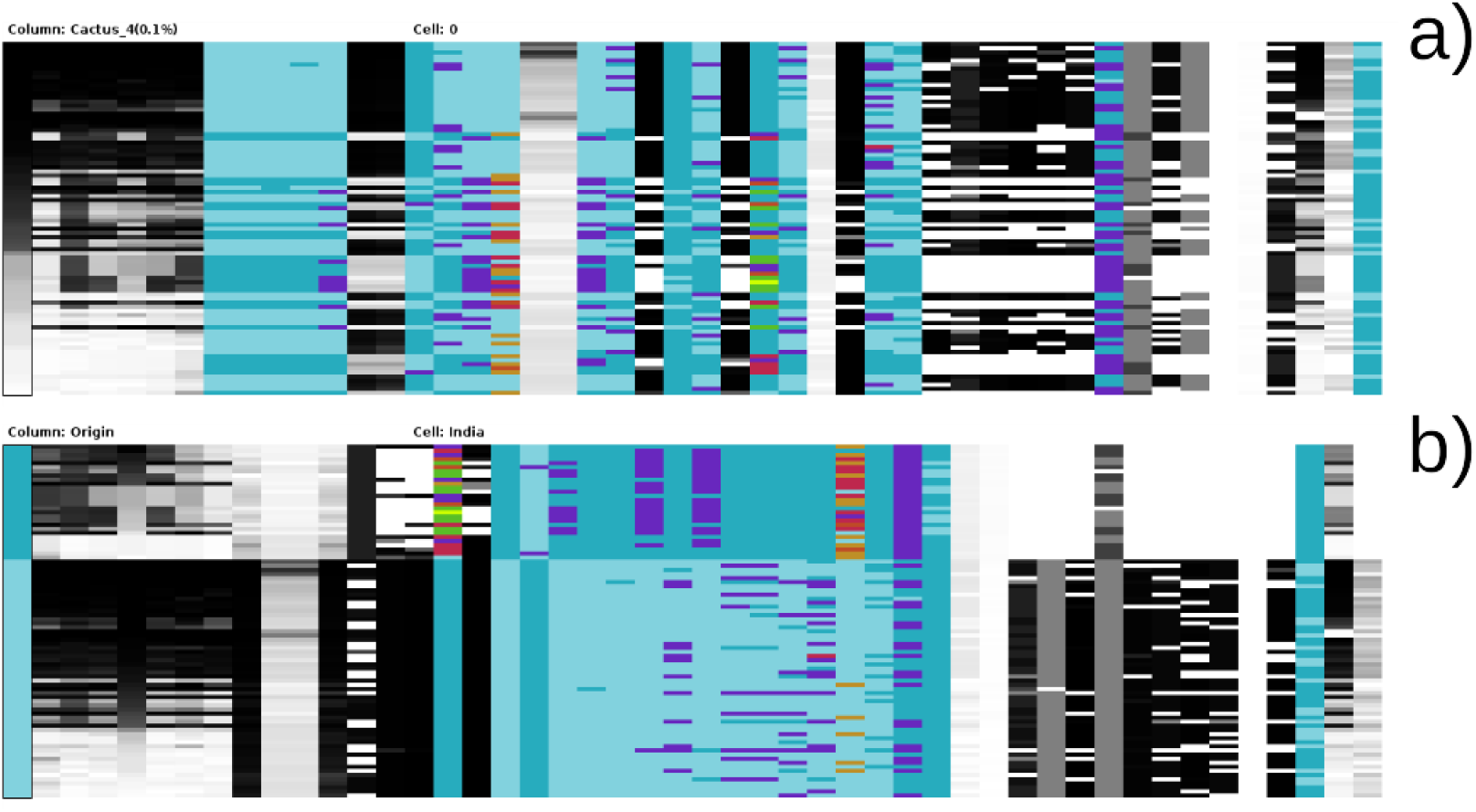
Column view in Mango. The column view is sorted by cactus 4 (a) and sorted by data origin (b). Numerical data is shown on a grey scale, e.g., columns 1 through 8, which contain values between 0 and 1. Text labels are coded into 28 colors, with colors reused as needed to accommodate more distinct labels. Hovering the mouse over a cell displays the column header and the content of the cell.

Mango also reveals clearer correlations, such as between read coverage and the number of reads. These variables also correlate with the sample origin, indicating possible differences in sequencing protocols that warrant further investigation.

Figure 2 shows the corresponding column view, sorted by cactus 4 (Figure 2a), and by sample origin (India or Norway, Figure 2b). This visualization presents individual elements organized into columns and rows. Numerical data is shown in a grey scale, scaled to the minimum and maximum values in each row. For text data, Mango assigns colors to each unique text category. For example, the sample source category “blood” is light blue, “urine” is purple, etc. The order of columns is based on the selected column in the correlation view, and all columns can be sorted by any column in the view, where numerical columns sorted numerically and text columns sorted alphanumerically.

**Figure 2.**
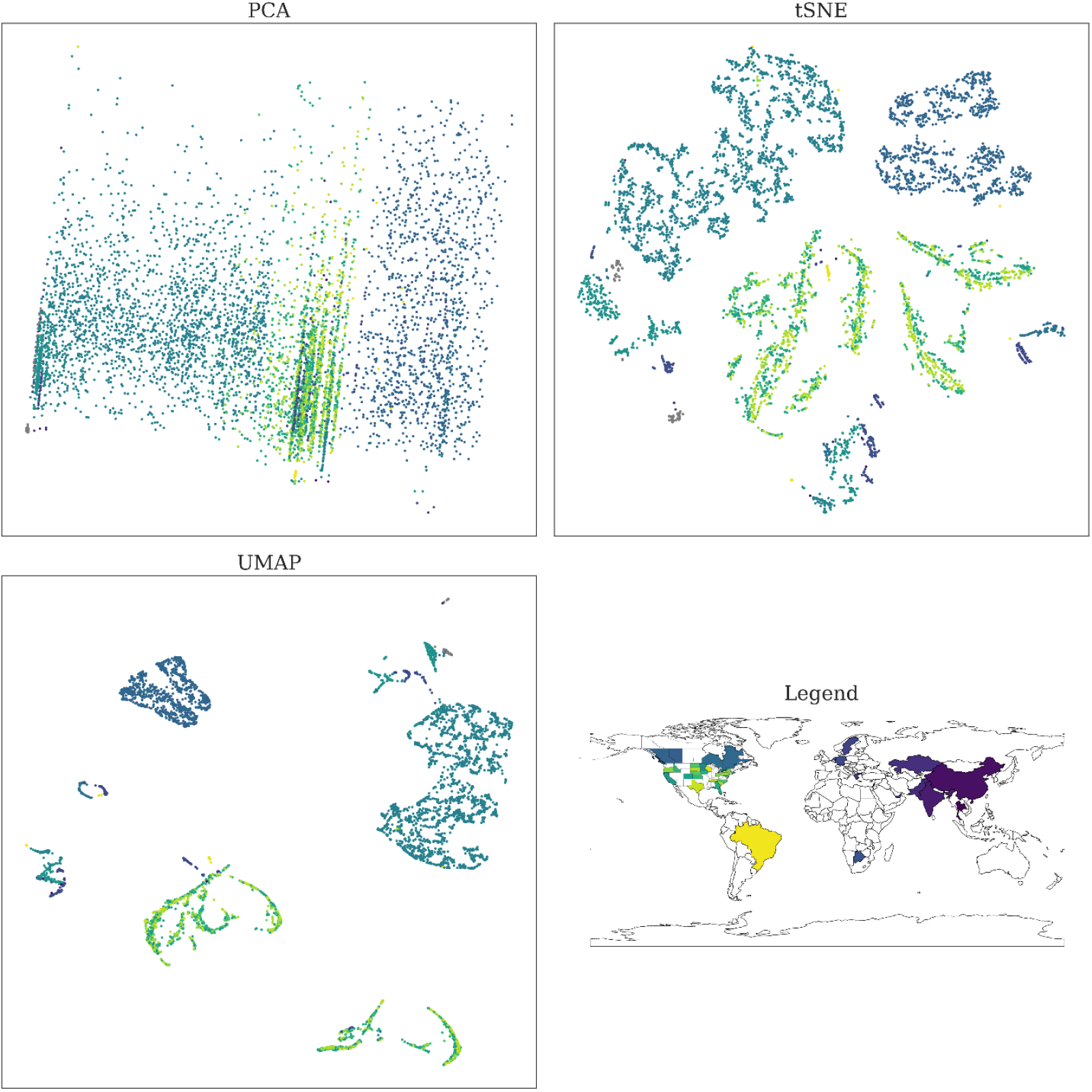
Dimensionality reduction plots of the NCBI and Avershina et al. datasets. Shown are PCA (a), tSNE (b), and UMAP (c). The NCBI dataset is colored by location (d). The distinct structure in the dimensionality reduction plots indicates that geographic location is a major factor influencing the resistance profile of *E. coli*.

Mango was also run on a larger dataset (the NCBI dataset) of publicly available *E. coli* isolates collected from various geographic locations, along with their phenotypic susceptibility or resistance to antibiotics. This dataset reveals a high correlation between the origin of the isolates and their phenotypic susceptibility (e.g. erythromycin, rho=0.78, p < 10^−5^), suggesting possible batch effects. For example, the resistance profile against erythromycin shows the highest correlation with “Location” (rho = 0.79). These trends identified by Mango are also evident in the structure of the dimensionality reduction plots, which are colored by the variable with the strongest correlation (Figure 3).

We next applied Mango to a larger RNA-Seq data set of fungal gene expression in Norway spruce roots, where one cohort was nutrient-enriched, and the control cohort was nutrient-limited (14). Almost instantly, Mango confirmed the key findings from the paper: the gene expression of fungi in the spruce varies significantly between the two groups but isn’t linked to any of the metadata (see right end of the plot). Figure 4 highlights the top correlations, with control samples shown in blue and fertilized samples in red.

**Figure 4.**
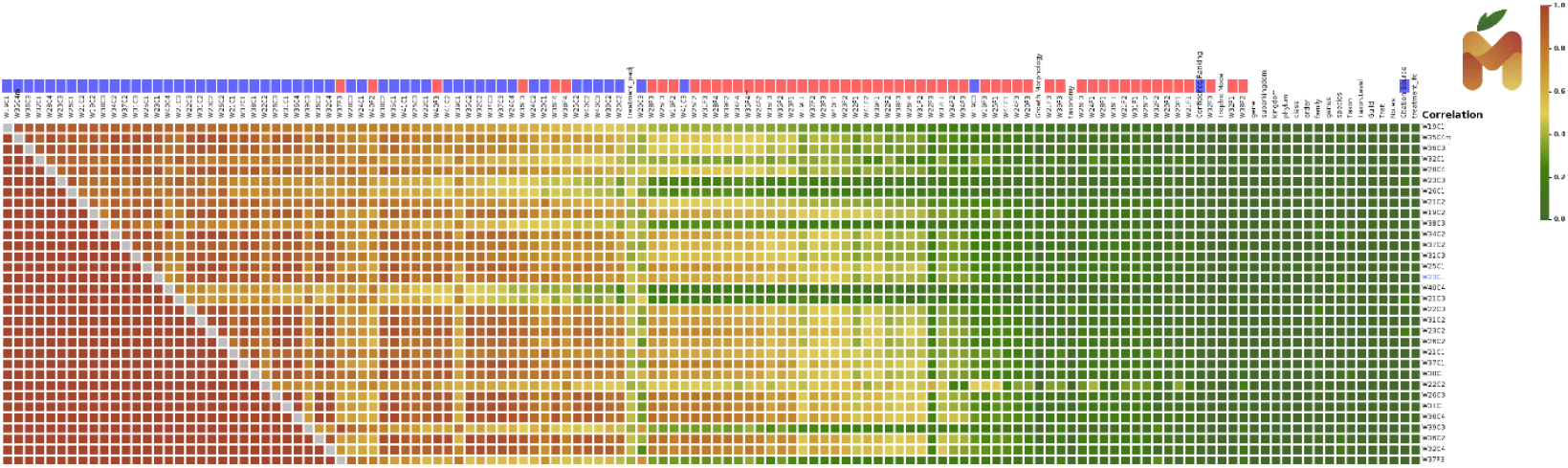
RNA-Seq expression of fungal populations in the roots of Norway spruce trees. One cohort, marked in blue, was nutrient-limited, while the other cohort, marked in red, was nutrient-enriched. The cohorts correlate with each other but are not related to any metadata (see right end of the plot).

## Discussion

Biological data sets are not only becoming larger, but also more complex and multi-modal. These data sets may aim to integrate data from different platforms, like genomics, transcriptomics, epigenetics, proteomics, spatial information, clinical data etc. In order to gain insight and discovery from these complex data, researchers increasingly rely on machine learning approaches which are able to find relationships and associations in these multi-dimensional data. The challenge, however, is ensuring that these analyses find meaningful associations rather than those caused by confounding factors (6). Given the scale and cost required to procure biological samples and omics data, optimal study design is often too costly, increasing the risk of batch effects (15). As demonstrated with an *E. coli* dataset (NCBI dataset), where genomic information was used to predict antibiotic resistance, the results in reality predict the location where the samples were collected. Mango’s interactive GUI makes it easy to identify such unwanted predictions with just a single mouse click, indicating that other analysis methods might be more effective for predicting antibiotic resistance from genotypes.

A popular graphical tool designed specifically for associating microbial genomic information to phenotypic traits is Scoary2 (16). This tool enables the analysis of tens of thousands of traits, and its data visualization features can facilitate the discovery of previously unknown bacterial genotype-phenotype connections. However, this tool has two main limitations that Mango addresses: (a) it violates the assumptions of Fisher’s test, meaning that resulting correlation values can only be interpreted as arbitrary scores; and (b) its pairwise comparison algorithm can only handle binary phenotypic events, not continuous or Brownian motion-type transitions, which limits the types of data that can be integrated. Continuous traits can be binarized, but this poses a risk of improper phenotypic classification or data being discarded when it does not fit a Gaussian mean, often requiring manual intervention and expertise to overcome.

Visualizing and exploring large data sets can be challenging due to the large number of possibilities in which data can be presented. Single numbers, such as numerical correlations, can also be misleading because they do not show the structure of the underlying data. Here, we introduce Mango, an interactive visualization and analysis tool that presents data at two levels: (a) correlations between data columns based on text-to-text, text-to-numeric, and numeric-to-numeric comparisons; (b) a more detailed column view, where each data point is accessible. Being able to compute correlations between heterogeneous data entries offers an efficient way to browse data systematically, while access to the most detailed level allows for deeper exploration. Additionally, Mango can serve as a pre-screening tool to estimate the strength of signals within the data, as demonstrated by the Norway spruce fungal gene expression dataset, which can motivate more in-depth analyses in early stages of research projects.

The user interface is designed to be interactive; for example, clicking on a data label in the correlation view sorts the entire matrix. To facilitate easier navigation through the two half-matrices, correlation, and p-value, hovering over a data point highlights the corresponding data point in both the correlation and p-value sections. The sorting of columns is consistent with the column view, where data can be sorted by each column. Each view can be exported as a Portable Network Graphics (PNG) file for presentation, sharing, and analysis.

Furthermore, it is important to emphasize that Mango is not limited to biomedical data sets and can be used for visualizing and exploring any data, as long as the data is organized in a tabular format.

## Conclusions

Mango is an interactive software tool for the quick analysis of mixed-type data sets, aiming at quickly and visually uncovering any biases in the data and to encourage more in-depth analyses early on in research projects if the results look promising. To the best of our knowledge, Mango is unique by its interactive user interface and by the focus on intuitive use, even by non-experts. Furthermore, Mango implements a novel algorithm for correlating categorical with numeric variables, regardless of distribution – Median-Ranked Label Encoding. This allows for direct comparison between numeric-numeric correlation, categorical-categorical correlation, and categorical-numeric correlation. We note that while we developed Mango for biological and medical data sets, it can be applied to any kind of tabular data.

## Methods

### Statistics Implementation

Mango aims to quantify the association between metadata variables. It does this by using correlation coefficients. For the association of numerical variables, Mango uses the non-parametric Pearson’s rank correlation coefficient (17) as follows. Given two numerical variables of size *n*

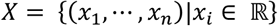

And

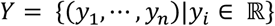

the variables are converted to ranks

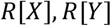

After which the correlation coefficient *r*_*s*_ is calculated using the sum of squared differences between ranks as

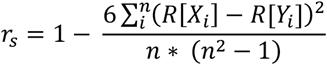

Correlation significance is tested using the Fisher transformation

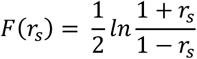

then the approximately normally distributed z-score

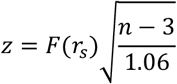

and a p-value can be computed using the standard normal distribution.

Mango quantifies the association between non-numeric variables using Cramér’s V method (18) as follows. Given two categorical variables

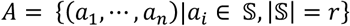

and

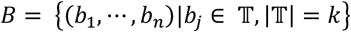

of sample size *n*, the chi-squared statistic is calculated by constructing a contingency table

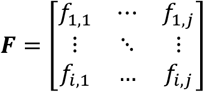

Where

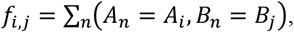

Then

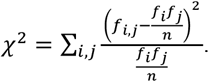

Using *χ*^2^, Cramér’s coefficient is computed as

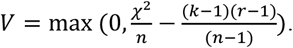

Since this coefficient tends to overestimate the strength of association, it is bias-corrected according to

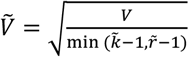

With

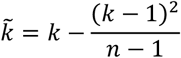

and

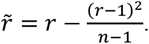

In addition, *χ*^2^ is used to compute a p-value using the cumulative chi-squared distribution with (*k* − 1)(*r* − 1) degrees of freedom using the lower incomplete gamma series expansion method.

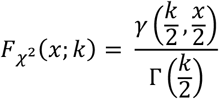

where the lower incomplete gamma function *γ* is computed via series expansion as

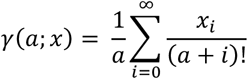

iteratively until convergence. The result is adjusted to account for the exponential decay in the tail of the incomplete gamma function as

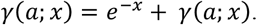

Associations between a numeric and non-numeric variable are calculated by first discretizing the numeric variable using what we call “Median-Ranked Label Encoding (MRLE)”. Given a numerical variable of size *n*

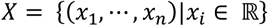

and a categorical variable of size *n*

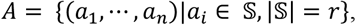

the numerical variable *X* is first grouped by category in *A* as follows

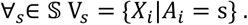

Next, the group-wise medians are calculated as

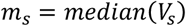

after which, the categories are sorted by their group medians. With a total order ≺ defined over 𝕊:

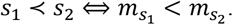

Finally, the original categorical labels are replaced with their corresponding median-sorted ranks. Let *rank*(*s*) be the position of *s* in the ordered list [*s*^(1)^, *s*^(2)^, …, *s*^(*k*)^], then:

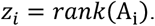

Then the MRLE-encoded categorical variable *A is*

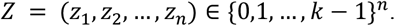

Once the categorical variable is encoded, the Spearman’s rank correlation coefficient and p-value are computed between *X* and *Z* as before.

After computing a correlation coefficient and a p-value for each pairwise combination of variables, Mango corrects the p-values for multiple comparisons using the Holm-Bonferroni method (19). Briefly, *m* p-values are sorted in ascending order, and then the adjusted p-value

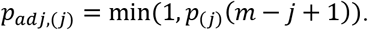

### Benchmarking data sets

The Avershina et al. (AMR) dataset includes genomic and minimum inhibitory concentration (MIC) data of 58 Norwegian and 28 Indian *Escherichia coli* isolates collected between 2015 and 2019 as described in the original publication (12). Variants were called using snippy (20) (v4.6.0) with default parameters. Subsequently, Saguaro (13) was used to generate cacti, modelling local phylogenetic homology without a priori hypotheses.

The NCBI dataset was constructed by selecting 7834 *E. coli* isolates with available MIC antibiotic susceptibility phenotypes from the NCBI isolates browser (21). Additional metadata was collected from the BioSample database using the Entrez utility, including geolocation and detailed antibiogram data. The data was filtered to remove sparse (>40% missing) and low information (>90% unique) variables. After filtering, the data contained 6538 isolates and 31 variables.

### Dimensionality Reduction

Numerical data were normalized, and categorical data were label-encoded using sklearn (22) (v. 1.3.2). Dimensionality reduction was performed using principal component analysis (PCA), and t-distributed stochastic neighbor embedding (t-SNE) implemented in sklearn, and uniform manifold approximation and projection (23) (UMAP).

## Abbreviations

GUI: graphical user interface
MIC: minimum inhibitory concentration.
PCA: principle component analysis
t-SNE: t-distributed stochastic neighbor embedding
UMAP: uniform manifold approximation and projection

## Declarations

### Ethics approval and consent to participate

Not applicable

### Consent for publication

Not applicable

### Availability of data and materials

Project name: Mango

Project home page: https://bitbucket.org/sverre-phd-work/mango/src/main/

Operating system: Linux

Programming language: C++, Java

Other requirements: openjdk >= 11.0

License: GNU GPL3

Any restrictions to use by non-academics: None other than specified in the license.

The datasets analyzed during the current study are available in the following repositories:

‐ Avrashina et al.dataset: https://www.ebi.ac.uk/ena/browser/view/PRJEB60478 and https://www.ncbi.nlm.nih.gov/bioproject/658789 (12)
‐ NCBI dataset: https://www.ncbi.nlm.nih.gov/pathogens/isolates/(21)
‐ Fungal gene expression dataset: enrichment tests can be accessed on the BRA web app (https://www.boreal-atlas.info/), raw RNA-seq data are available in the European nucleotide archive (ENA), https://www.ebi.ac.uk/ena/browser/ (accession no. PRJEB35805) (14).

### Authors’ contributions

RA, SB, and MGG conceived the study. SB designed and MGG implemented the GUI. SB and MGG designed and carried out the statistical analyses. All authors jointly wrote, edited, and approved the manuscript.

## Acknowledgements

Not applicable

## References

1. Yu Y, Mai Y, Zheng Y, Shi L. Assessing and mitigating batch effects in large-scale omics studies. Genome Biol. 2024 Oct 3;25(1):254.

2. Ardila CM, González-Arroyave D, Tobón S. Machine learning for predicting antimicrobial resistance in critical and high-priority pathogens: A systematic review considering antimicrobial susceptibility tests in real-world healthcare settings. PLOS One. 2025 Feb 25;20(2):e0319460.

3. Sakagianni A, Koufopoulou C, Feretzakis G, Kalles D, Verykios VS, Myrianthefs P, et al. Using Machine Learning to Predict Antimicrobial Resistance―A Literature Review. Antibiotics. 2023 Mar;12(3):452.

4. Li Y, Cui X, Yang X, Liu G. Artificial intelligence in predicting pathogenic microorganisms’ antimicrobial resistance: challenges, progress, and prospects. Front Cell Infect Microbiol [Internet]. 2024 Nov 1 [cited 2025 Sept 9];14. Available from: https://www.frontiersin.org/journals/cellular-and-infection-microbiology/articles/10.3389/fcimb.2024.1482186/full

5. Earle SG, Wu CH, Charlesworth J, Stoesser N, Gordon NC, Walker TM, et al. Identifying lineage effects when controlling for population structure improves power in bacterial association studies. Nat Microbiol. 2016 Apr 4;1(5):1–8.

6. O’Donoghue SI. Grand Challenges in Bioinformatics Data Visualization. Front Bioinforma [Internet]. 2021 June 17 [cited 2025 June 12];1. Available from: https://www.frontiersin.org/journals/bioinformatics/articles/10.3389/fbinf.2021.669186/full

7. Stajich JE, Block D, Boulez K, Brenner SE, Chervitz SA, Dagdigian C, et al. The Bioperl Toolkit: Perl Modules for the Life Sciences. Genome Res. 2002 Oct 1;12(10):1611–8.

8. Gentleman RC, Carey VJ, Bates DM, Bolstad B, Dettling M, Dudoit S, et al. Bioconductor: open software development for computational biology and bioinformatics. Genome Biol. 2004 Sept 15;5(10):R80.

9. Grüning B, Dale R, Sjödin A, Chapman BA, Rowe J, Tomkins-Tinch CH, et al. Bioconda: sustainable and comprehensive software distribution for the life sciences. Nat Methods. 2018 July;15(7):475–6.

10. Chelaru F, Smith L, Goldstein N, Bravo HC. Epiviz: interactive visual analytics for functional genomics data. Nat Methods. 2014 Sept;11(9):938–40.

11. Sievert C. Interactive Web-Based Data Visualization with R, plotly, and shiny. New York: Chapman and Hall/CRC; 2020. 470 p.

12. Avershina E, Sharma P, Taxt AM, Singh H, Frye SA, Paul K, et al. AMR-Diag: Neural network based genotype-to-phenotype prediction of resistance towards \upbeta-lactams in Escherichia coli and Klebsiella pneumoniae. Comput Struct Biotechnol J. 2021;19:1896–906.

13. Zamani N, Russell P, Lantz H, Hoeppner MP, Meadows JR, Vijay N, et al. Unsupervised genome-wide recognition of local relationship patterns. BMC Genomics. 2013 May 24;14(1):347.

14. Law SR, Serrano AR, Daguerre Y, Sundh J, Schneider AN, Stangl ZR, et al. Metatranscriptomics captures dynamic shifts in mycorrhizal coordination in boreal forests. Proc Natl Acad Sci. 2022 June 28;119(26):e2118852119.

15. Tom JA, Reeder J, Forrest WF, Graham RR, Hunkapiller J, Behrens TW, et al. Identifying and mitigating batch effects in whole genome sequencing data. BMC Bioinformatics. 2017 July 24;18(1):351.

16. Roder T, Pimentel G, Fuchsmann P, Stern MT, von Ah U, Vergères G, et al. Scoary2: rapid association of phenotypic multi-omics data with microbial pan-genomes. Genome Biol. 2024 Apr 11;25(1):93.

17. Spearman C. The Proof and Measurement of Association between Two Things. Am J Psychol. 1904 Jan;15(1):72.

18. Cramér H. Mathematical methods of statistics. Princeton: Princeton university press; 1991. (Princeton mathematical series).

19. Chen SY, Feng Z, Yi X. A general introduction to adjustment for multiple comparisons. J Thorac Dis. 2017 June;9(6):1725–9.

20. Seemann T. tseemann/snippy [Internet]. 2025 [cited 2025 May 12]. Available from: https://github.com/tseemann/snippy

21. National Center for Biotechnology Information, National Library of Medicine, National Institutes of Health, Bethesda, MD 20894, USA. The NCBI Pathogen Detection Project. [Internet]. 2016 [cited 2025 May 12]. Available from: https://www.ncbi.nlm.nih.gov/pathogens/

22. Pedregosa F, Varoquaux G, Gramfort A, Michel V, Thirion B, Grisel O, et al. Scikit-learn: Machine Learning in Python. J Mach Learn Res. 2011;12(85):2825–30.

23. McInnes L, Healy J, Melville J. UMAP: Uniform Manifold Approximation and Projection for Dimension Reduction [Internet]. arXiv; 2020 [cited 2025 May 12]. Available from: http://arxiv.org/abs/1802.03426

